# The HSV-1 LAT enhancer regulates locus-specific H3.1 enrichment during early lytic infection

**DOI:** 10.64898/2026.09.30.755688

**Authors:** Kayley A. Manuel, Tori A. Fraser, Mason A. Shipley, Donna M. Neumann

## Abstract

The Herpes Simplex Virus 1 (HSV-1) genome undergoes extensive chromatinization following entry into the nucleus, yet how viral regulatory elements influence this process during lytic infection remains poorly understood. The HSV-1 latency-associated transcript enhancer (LTE) is well established as a key element in the regulation of latency and reactivation, but its role during the productive infection has not been defined. Here, we identify a role for the LTE in regulating chromatin composition on the lytic HSV-1 genome. Deletion of a 310-bp region within the LTE selectively reduced enrichment of the canonical histone H3.1 at specific viral lytic genes, including ICP4 and ICP8, without altering H3.3 enrichment. This phenotype was not reproduced using a second LTE deletion that disrupted enhancer activity, indicating that regulation of H3.1 enrichment is separable from the function of the enhancer. LTE-dependent H3.1 enrichment persisted in the absence of viral DNA replication and was accompanied by reduced occupancy of the H3.1 histone chaperone CAF-1. However, depletion of CAF-1 did not substantially reduce H3.1 enrichment, indicating that additional mechanisms contribute to H3.1 association with the viral genome. Together, these findings reveal that the HSV-1 LTE regulates locus-specific chromatin composition during early lytic infection and identify a previously unrecognized connection between a viral enhancer and histone H3.1 enrichment on the incoming viral genome.

**Author Summary:** Viruses must organize their genetic material inside infected cells to control when viral genes are turned on and off. After HSV-1 enters a cell, its DNA becomes associated with histones. Although HSV-1 contains regulatory sequences that control viral gene expression, it is not well understood how these sequences influence the chromatin deposited on the viral genome during infection. In this study, we discovered that the HSV-1 enhancer (LTE), a viral DNA element previously known primarily for its role in latency and reactivation, also regulates the chromatin landscape during lytic infection. Removing a specific region of this enhancer reduced the association of H3.1 early after infection. This effect was also associated with decreased binding of the cellular histone-loading complex CAF-1. These changes in the chromatin landscape were accompanied by altered expression of specific viral genes. Together, our findings reveal an unexpected role for the LTE in regulating the chromatin environment of the viral genome during lytic infection and provide new insight into how viral DNA elements interact with cellular chromatin machinery to control infection.

## Introduction

Herpes Simplex Virus Type 1 (HSV-1) is a double-stranded DNA virus that causes lifelong infection in ∼2/3 of the global population [1]. Disease manifestation ranges from mild conditions, such as the development of cold sores to the more severe keratitis and encephalitis [1–4]. HSV-1 has a biphasic life cycle characterized by three distinct infection phases: primary (lytic) infection, latency, and reactivation. Lytic infection begins in epithelial cells, where the virus delivers its genome to the host nucleus and initiates a temporal gene expression cascade to produce viral proteins, followed by viral DNA replication and ultimately assembly of infectious virions. These progeny virions then undergo retrograde axonal transport to sensory neurons where the virus establishes latency, characterized by the repression of nearly all viral genes. HSV-1 can undergo periodic reactivation induced by various environmental stressors, allowing the virus to travel via axons back to the epithelium and reestablish a productive infection. These distinct infection stages are largely controlled by epigenetic mechanisms that regulate accessibility and transcriptional activity of the viral genome. Although considerable progress has been made in defining the chromatin mechanisms that control HSV-1 latency and reactivation, how specific elements within the viral genome regulate chromatin organization during the initiation of lytic infection remains poorly understood.

Upon entry into the host nucleus, the HSV-1 genome rapidly associates with nucleosomes as early as 1 hpi. Unlike cellular chromosomes, the incoming HSV-1 genome initially acquires chromatin in an irregular and destabilized manner [5, 6]. Histone deposition therefore represents an early event that may influence the transcriptional state of the HSV-1 genome. Histone occupancy increases and peaks at 3 hpi, near the beginning of viral DNA replication [6, 7], and multiple studies have shown that HSV-1 utilizes histone mobilization to contribute to the regulation of gene expression during lytic infection [7–10]. Importantly, histone deposition on the viral genome occurs before viral DNA replication and does not require viral protein synthesis [8, 10], suggesting that establishment of viral chromatin is an early regulatory event rather than a consequence of viral replication. However, the mechanisms that determine the composition and distribution of histones at specific viral loci remain under established.

Histone variants may provide one mechanism through which distinct chromatin environments are established on viral genomes. Histone H3 exists primarily as the canonical H3.1 variant and the replication independent H3.3 variant, both of which are deposited through distinct histone chaperone pathways. For example, H3.1 is expressed during S phase and is classified as a replication-dependent histone that is predominantly loaded by the chromatin assembly factor 1 (CAF-1) [11–13]. On the other hand, H3.3 is replication-independent and mainly loaded by the histone regulator A complex (HIRA) [11]. Despite differing in amino acid sequence by 5 residues [13], H3.1 and H3.3 have distinct distributions on genomes and are often associated with distinct chromatin environments. For example, H3.1 is frequently associated with heterochromatic regions of genomes and, in part, represses transcription [11, 12] and is mobilized onto the viral genome prior to DNA replication [9, 10]. On the other hand, H3.3 is associated with euchromatic regions that are readily accessible for transcription [14, 15]. Further, H3.3 enrichment is found at active gene promoters, active gene bodies, and enhancers [16] and H3.3 has been shown to associate with viral gene promoters throughout lytic infection to support gene expression [7]. These distinct functions raise the possibility that specific viral regulatory elements influence the incorporation of individual histone variants at specific genomic loci.

Enhancers are well-established regulators of cellular chromatin organization and gene expression through functioning as recruiters of transcription factors, influencing chromatin remodeling, and function over long genomic distances [17–19]. HSV-1 encodes a viral enhancer located in the 5’ exon region of the Latency Associated Transcript - a highly dynamic long non-coding RNA that has roles in chromatin organization, neuronal protection, the establishment of latency and in efficient reactivation [20–26]. The LTE can enhance transcription of nearby and distal genes, and we have recently shown that the LTE plays a role in establishing long-range chromatin architecture during latency [27–29]. Thus, the LTE represents a potential viral regulatory element capable of coordinating chromatin organization and transcription across the HSV-1 genome.

The HSV-1 LTE has been studied extensively for its roles in latency and reactivation, however its potential function in regulating the viral genome during the productive infection remains poorly understood. Previous studies utilizing two HSV-1 recombinant viruses with partial deletions of the LTE, including 17Δ348 and 17ΔA (**Fig. 1**), established important roles for the enhancer in establishment and maintenance of latency, epigenetic regulation of latency, and efficient reactivation [27, 29–31]. Of note, while these recombinant viruses have been shown to have attenuated reactivation phenotypes, we found no measurable lytic growth defect [27, 31], suggesting that the LTE may have additional functions during the lytic infection that are distinct from its already established roles in latency and reactivation. Whether the LTE influences the histone deposition on incoming HSV-1 genomes during the establishment of lytic infection has not yet been determined.

**Figure 1:**
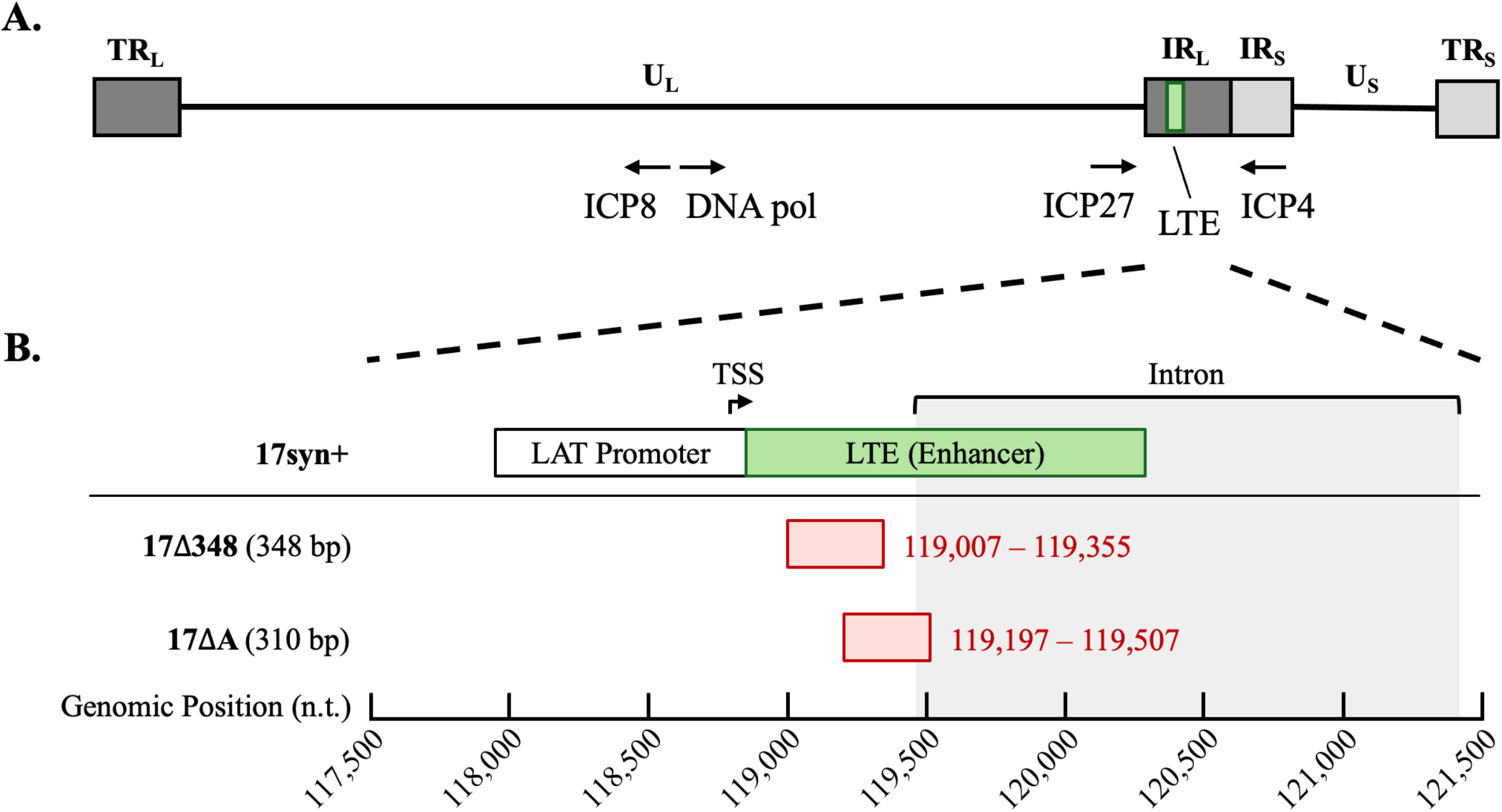
Map of WT HSV-1 (17syn+) and LTE deleted viruses (17Δ348 and 17ΔA). **A)** Linear map of the HSV-1 genome. Genes referred to in the text are shown with their relative genomic locations. Note: LTE and ICP4 are located in repeat regions, the figure shows only the internal repeat copies. **B)** Map of enhancer deleted viruses 17Δ348 and 17ΔA compared to 17syn+.

Here, we identify a previously unrecognized role for the HSV-1 LTE in regulating locus-specific enrichment of H3.1 during early lytic infection. Using the two independent partial LTE deletion viruses described above, we found that deletion of a specific 310 bp LTE region selectively reduced H3.1 enrichment at specific loci including the immediate early gene ICP4, without reducing H3.3 enrichment. LTE-dependent H3.1 enrichment at ICP4 occurred before viral DNA replication and was accompanied by reduced occupancy of the canonical H3.1 histone chaperone CAF-1. However, depletion of CAF-1 did not substantially impair H3.1 enrichment, indicating that CAF-1 contributed to, but was not solely responsible for, H3.1 deposition on HSV-1 lytic genes. Together, these findings revealed that the LTE can selectively coordinate histone variant enrichment and chromatin assembly at specific loci on the incoming HSV-1 genome, providing a mechanism through which HSV-1 regulates its chromatin landscape during the productive infection.

## Results

### Deletion of a 310 bp LTE region selectively reduces H3.1 enrichment at specific HSV-1 lytic genes

To investigate whether LTE influenced occupancy of the canonical histone, H3.1, we utilized two partial LTE deletion viruses and their rescues, 17ΔA and 17Δ348, along with the wild type (WT) 17syn+ (**Fig. 1**). To quantify histone enrichment, we performed chromatin immunoprecipitation coupled with quantitative PCR (ChIP-qPCR) using an antibody specific to H3.1. Normal Human Dermal Fibroblast (NHDF) cells were infected at an MOI of 1 with respective viruses and samples were collected at 2 hpi to quantify H3.1 occupancy at various lytic genes. We selected the timepoint of 2 hpi to examine histone deposition during the early phase of lytic infection, prior to substantial viral DNA replication. H3.1 enrichment was validated using the cellular GAPDH locus as a positive control in ChIP. Only samples with greater than 3-fold enrichments of H3.1 on GAPDH compared to the negative control antibody, IgG, were considered validated and used for further analyses (**Fig. S1**). Next, we quantified H3.1 enrichment on IE genes (ICP4 and ICP27) and E genes (ICP8 and DNA polymerase). In all samples, we found that all four genes were significantly enriched with H3.1 at 2 hpi compared to IgG. Interestingly, when we assessed 17ΔA, we found that deletion of the 310 bp LTE region resulted in a significant decrease of H3.1 occupancy on ICP4 and ICP8 compared to both WT and rescue viruses (**Fig. 2A&C**). Additionally, the decrease in H3.1 occupancy was confined to only ICP4 and ICP8, as H3.1 occupancy was not significantly different on viral genes ICP27 and DNA polymerase in any of the viruses analyzed at this time point (**Fig. 2B&D**). These findings indicate that the LTE contributes to H3.1 enrichment at specific viral loci during early lytic infection.

**Figure 2:**
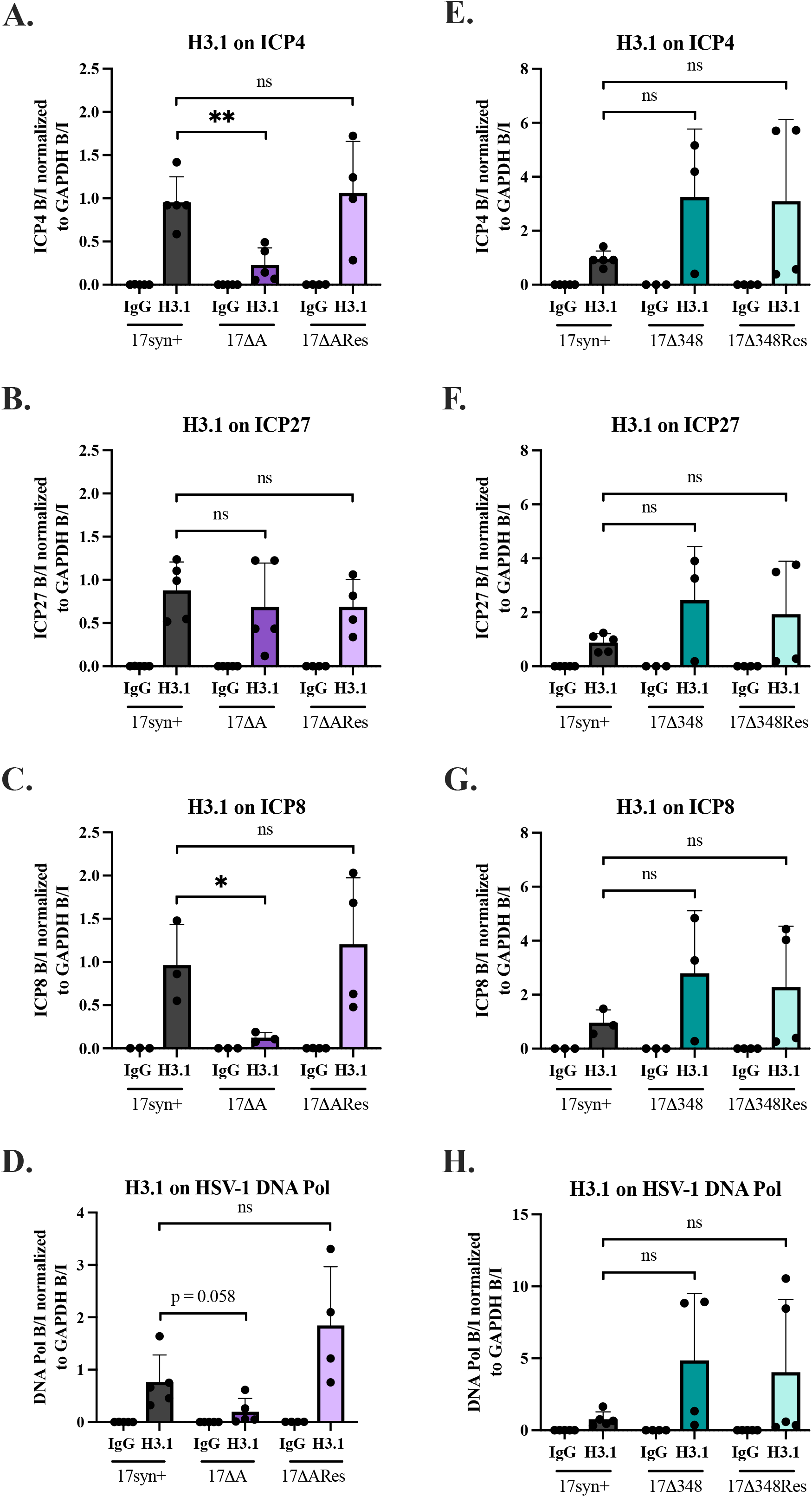
H3.1 enrichment on viral lytic genes in 17syn+, 17ΔA, 17ΔARescue, 17Δ348, and 17Δ348Rescue at 2 hpi. NHDF cells were infected with respective viruses at a MOI = 1, then ChIP-qPCR was performed to quantify H3.1 enrichment on viral lytic genes and normalized to H3.1 enrichment on the cellular control gene, GAPDH. H3.1 enrichment is plotted compared the non-specific binding control IgG (IgG values are the bound/input values of the respective viral gene). H3.1 enrichment compared to 17syn+ is shown for 17ΔA and 17ΔARescue in Figures 2A-D and 17Δ348 and 17Δ348Rescue in Figures 2E-H. The order of viral genes is as follows (top to bottom): ICP4, ICP27, ICP8, and HSV-1 DNA polymerase. N = 3-5. Unpaired two-tailed Student’s t-tests were performed to determine statistical significance between samples, means and standard deviations are plotted, * = p < 0.05, ** = p < 0.01, ns = not significant, p ≥ 0.05.

### A second LTE deletion that disrupts enhancer activity does not alter H3.1 enrichment

We next asked whether the effect of the LTE on H3.1 enrichment reflected disruption of its established enhancer activity or instead involved a distinct function of the deleted sequence. To address this question, we utilized an additional recombinant, 17Δ348 and its rescue to quantify H3.1 occupancy at the same viral lytic genes. The LTE deletion in 17Δ348 overlaps the 17ΔA deletion by 158 bp and contains an additional 190 bp of non-overlapping sequence that is also deleted (**Fig. 1B**). Both viruses have been shown to disrupt enhancer function [27, 28, 31]. Interestingly, we found that H3.1 enrichment on 17Δ348 was not significantly altered compared to WT or the 17Δ348 rescue virus at any of the lytic genes assayed (**Fig. 2E–H**). Thus, the reduction in H3.1 enrichment is not a general consequence of disrupting LTE enhancer activity but is instead associated with sequence(s) within the 310 bp region deleted in 17ΔA.

### Deletion of the LTE selectively reduces H3.1 enrichment without altering H3.3 variant enrichment

Having established that the 17ΔA deletion selectively alters H3.1 enrichment at specific viral loci, we next asked whether LTE deletion similarly affects enrichment of the replication-independent H3.3 histone variant. To determine whether the LTE globally regulates the enrichment of H3, we used ChIP-qPCR to quantitate H3.3 at the same viral loci in WT and 17ΔA. In contrast to H3.1, partial deletion of the LTE did not significantly affect H3.3 enrichment at any of the viral loci examined (**Fig. 3A–D).** Thus, the 17ΔA deletion selectively affects H3.1 enrichment without detectably altering H3.3 enrichment at the viral loci examined.

**Figure 3:**
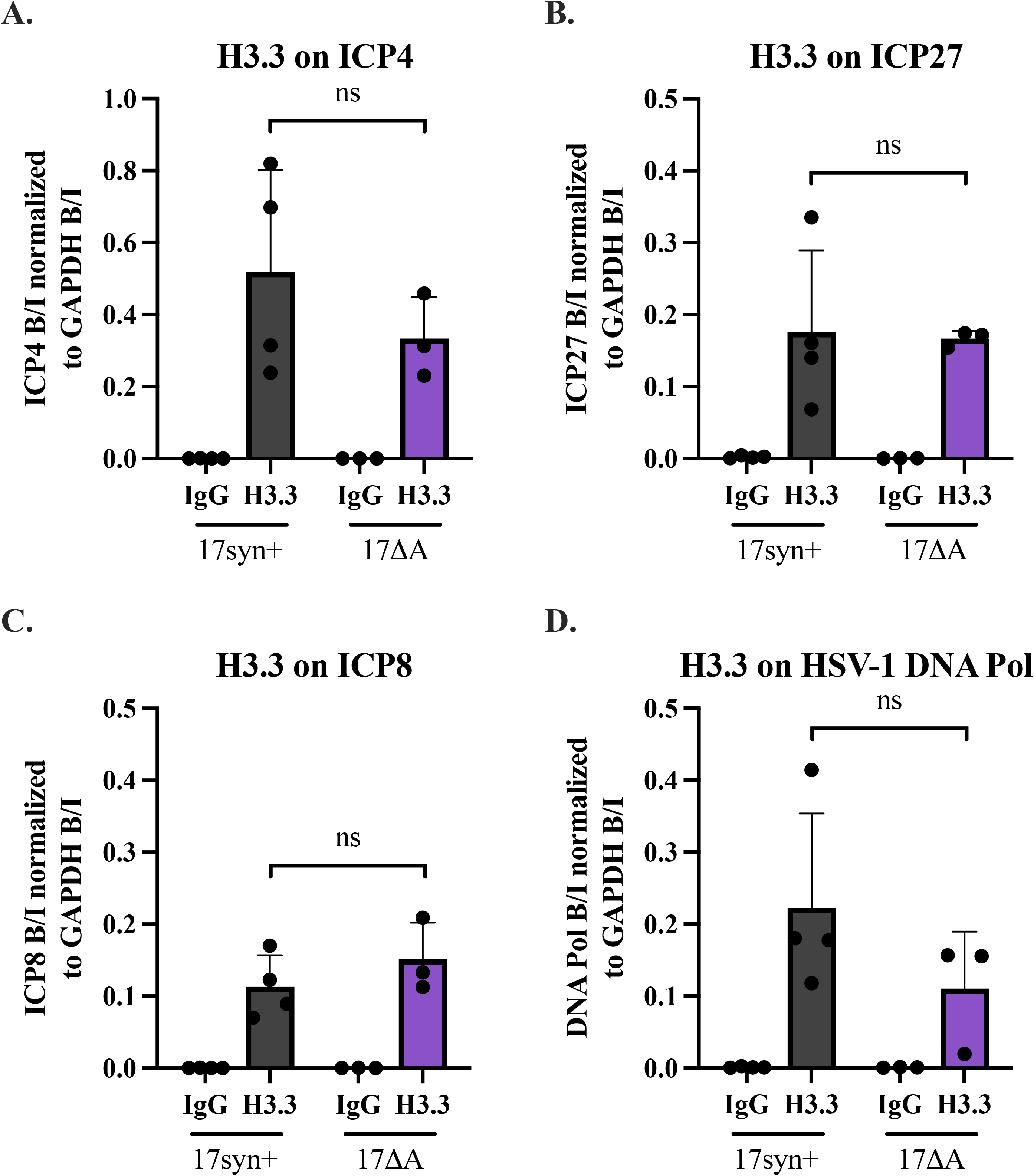
Histone 3.3 enrichment on viral lytic genes in 17syn+ and 17ΔA at 2hpi. NHDF cells were infected with respective viruses at a MOI = 1, then ChIP-qPCR was performed to quantify H3.3 enrichment on viral lytic genes and normalized to H3.3 enrichment on the cellular control gene, GAPDH. H3.3 enrichment is plotted against the non-specific binding control IgG (IgG values are the bound/input values of the respective viral gene). Viral genes: **A)** ICP4, **B)** ICP27, **C)** ICP8, and **D)** HSV-1 DNA polymerase. N = 3-4. Unpaired two-tailed Student’s t-tests were performed to determine statistical significance between samples, means and standard deviations are plotted, ns = not significant, p ≥ 0.05.

### LTE deletion alters viral gene expression during lytic infection

Because the LTE deletion selectively altered H3.1 enrichment at specific viral loci, we next asked whether this chromatin phenotype was associated with changes in viral gene expression during lytic infection. To determine this, we infected NHDF cells with WT, 17ΔA, or 17ΔA rescue viruses at an MOI of 1 and quantified viral transcripts by RT-qPCR (**Table S1)**. We examined the expression of the IE genes ICP4 and ICP27 and the E genes ICP8 and DNA polymerase over the course of infection. Viral transcript abundance increased over time for all viruses, consistent with the expected progression of the lytic transcriptional cascade. However, 17ΔA-infected cells exhibited reduced expression of several viral genes compared with WT-infected cells, including ICP27, ICP8, and DNA polymerase (**Fig. 4A&B**). Importantly, expression of these genes was restored toward WT levels in cells infected with the 17ΔA rescue virus (**Fig. 4C**), supporting a specific effect of the LTE deletion on viral gene expression. In contrast, ICP4 expression followed a distinct pattern and was not reduced in 17ΔA-infected cells. To compare viral gene expression at a later stage of infection, transcript abundance was assessed at 24 hpi. At this time point, ICP27, ICP8, and DNA polymerase transcripts remained reduced in 17ΔA compared with WT, whereas ICP4 transcript abundance increased in 17ΔA (**Fig. 4D–G**). Thus, deletion of the LTE that reduces H3.1 enrichment at specific viral loci is also associated with altered expression of several genes during the lytic infection. To determine whether these differences in viral transcript abundance were a consequence of altered viral replication, we performed single-cycle growth analyses in NHDF cells infected at an MOI of 1 and quantified viral genome copies and infectious virus production. 17ΔA did not exhibit a significant replication defect compared with WT, although a modest increase in viral genome copies and infectious virus production was observed at 8 hpi that was no longer apparent at 24 hpi or later (**Fig. S2**). These findings indicate that the altered viral transcript abundance observed following LTE deletion is not explained by a reduction in viral replication.

**Figure 4:**
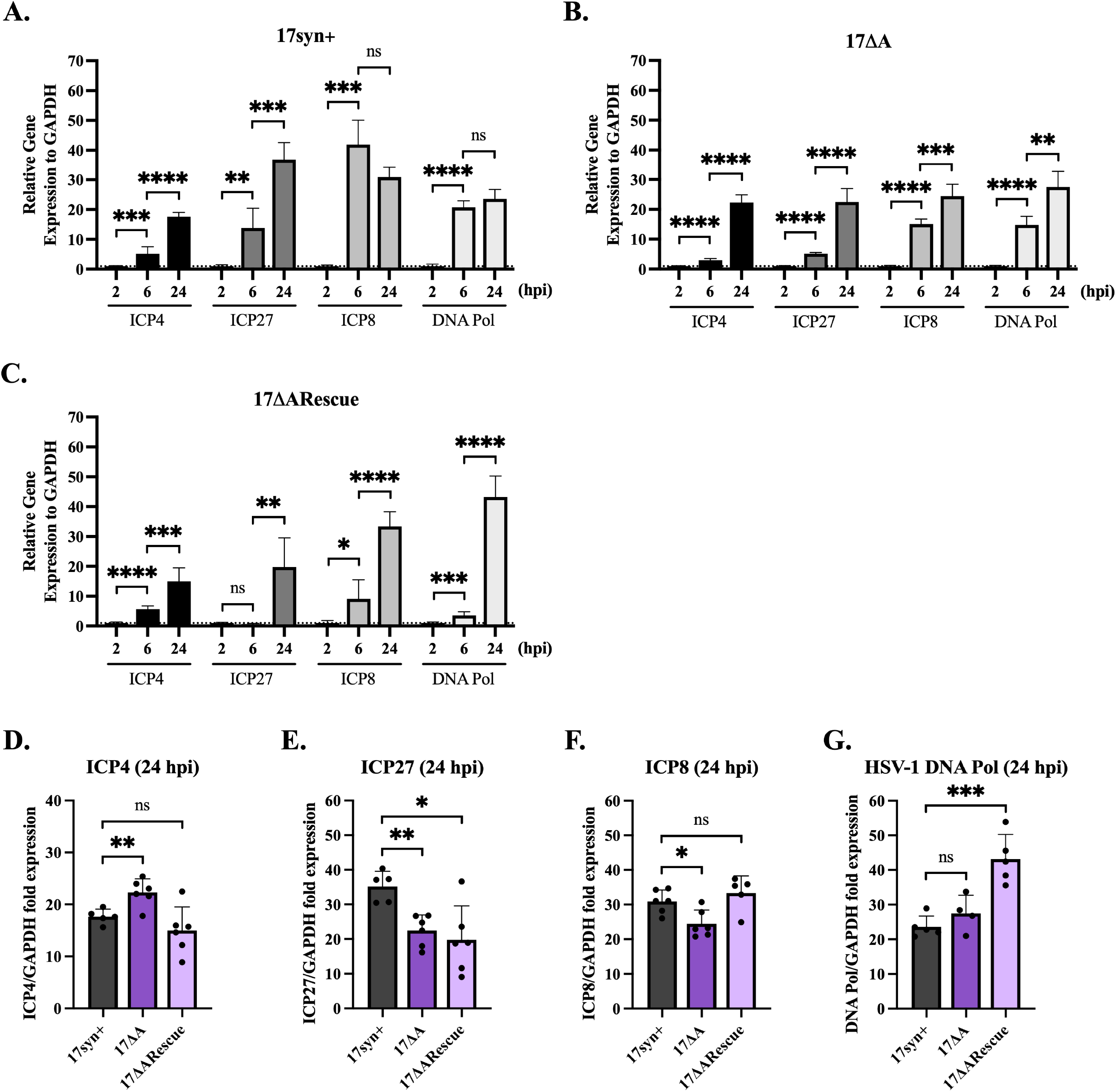
Gene Expression of 17syn+, 17ΔA, and 17ΔARescue at 2, 6, and 24 hpi. NHDF cells were infected with respective viruses at a MOI = 1, then qRT-PCR was used to quantify viral gene expression and normalized to the cellular control, GAPDH. To plot fold expression, respective genes were normalized to the 2 hpi sample. **A)** 17syn+ gene expression. **B)** 17ΔA gene expression. **C)** 17ΔARescue gene expression. N = 4-6. Data points from Graphs 4A-C were replotted to compare respective genes across all viruses at 24 hpi. Viral genes: **D)** ICP4, **E)** ICP27, **F)** ICP8, and **G)** HSV-1 DNA polymerase. Unpaired two-tailed Student’s t-tests were performed to determine statistical significance between samples, means and standard deviations are plotted, * = p < 0.05, ** = p < 0.01, *** = p < 0.001, **** = p < 0.0001, ns = not significant, p ≥ 0.05.

### LTE-dependent H3.1 enrichment at ICP4 occurs independently of viral DNA replication

We next asked whether LTE-dependent H3.1 enrichment required viral DNA replication. Although H3.1 is classically described as a replication-dependent histone in human cells, we investigated whether blocking replication would alter the H3.1 profile of 17ΔA. To do this, we lytically infected NHDF cells in the presence of the viral DNA replication inhibitor Phosphonoacetic acid (PAA). PAA is a pyrophosphate analog and competitively inhibits the HSV-1 DNA polymerase by binding to it to block chain elongation [32, 33]. Under the replication-inhibited conditions, 17ΔA again exhibited reduced H3.1 enrichment at ICP4 compared with both WT and 17ΔARescue viruses (**Fig. 5**). These findings demonstrate that LTE-dependent H3.1 enrichment at ICP4 occurs independently of viral DNA replication.

**Figure 5:**
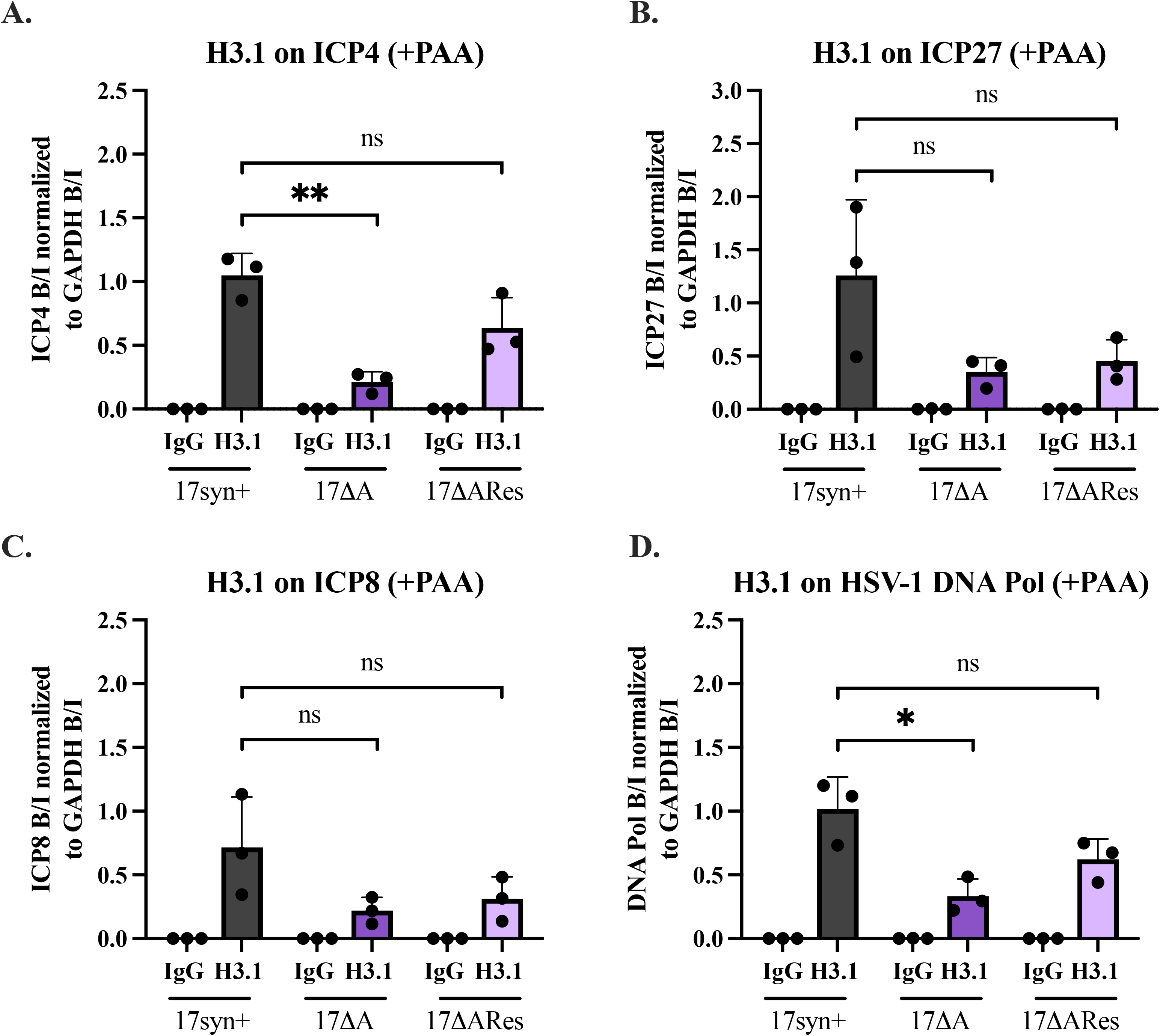
H3.1 enrichment on viral lytic genes in 17syn+, 17ΔA, 17ΔARescue at 2 hpi in the absence of viral replication (PAA inhibition). NHDF cells were infected with respective viruses at a MOI = 1 in the presence of PAA to block viral replication. Then, ChIP-qPCR was performed to quantify H3.1 enrichment on viral lytic genes and normalized to H3.1 enrichment on the cellular control gene, GAPDH. H3.1 enrichment is plotted against the non-specific binding control IgG (IgG values are the bound/input values of the respective viral gene). Viral genes: **A)** ICP4, **B)** ICP27, **C)** ICP8, and **D)** HSV-1 DNA polymerase. N = 3. Unpaired two-tailed Student’s t-tests were performed to determine statistical significance between samples, means and standard deviations are plotted, * = p < 0.05, ** = p < 0.01, ns = not significant, p ≥ 0.05.

### Deletion of LTE reduces CAF-1 occupancy at viral loci with reduced H3.1 enrichment

Having established that LTE deletion selectively reduces H3.1 enrichment at specific viral loci independently of viral DNA replication, we next asked whether this phenotype was associated with altered occupancy of the H3.1 histone chaperone chromatin assembly factor 1 (CAF-1). CAF-1 is a major histone chaperone for H3.1 and has been implicated in both replication-dependent and replication-independent chromatin assembly [13, 34]. The CAF-1 complex consists of three subunits: the large subunit p150 (CHAF1A), the middle subunit p60 (CHAF1B), and the small subunit p48 (RbAp48) [13]. Therefore, to address this question, we quantified occupancy of CAF-1 at the same viral genes that exhibited decreased H3.1 enrichment in 17ΔA during lytic infection. To assess CAF-1 occupancy, we performed ChIP-qPCR using an antibody directed against the large CAF-1 subunit, Chaf1a. CAF-1 enrichment at viral lytic genes was normalized to enrichment at the cellular GAPDH locus. We observed a significant decrease in CAF-1 occupancy at ICP4 and a corresponding decrease at ICP8 by 2 hpi in 17ΔA-infected cells compared with WT-infected cells (**Fig. 6A-D**). Thus, deletion of the LTE reduced CAF-1 occupancy at viral loci that also exhibited reduced H3.1 enrichment, suggesting that the LTE influenced the association of CAF-1 with specific regions of the viral genome.

**Figure 6:**
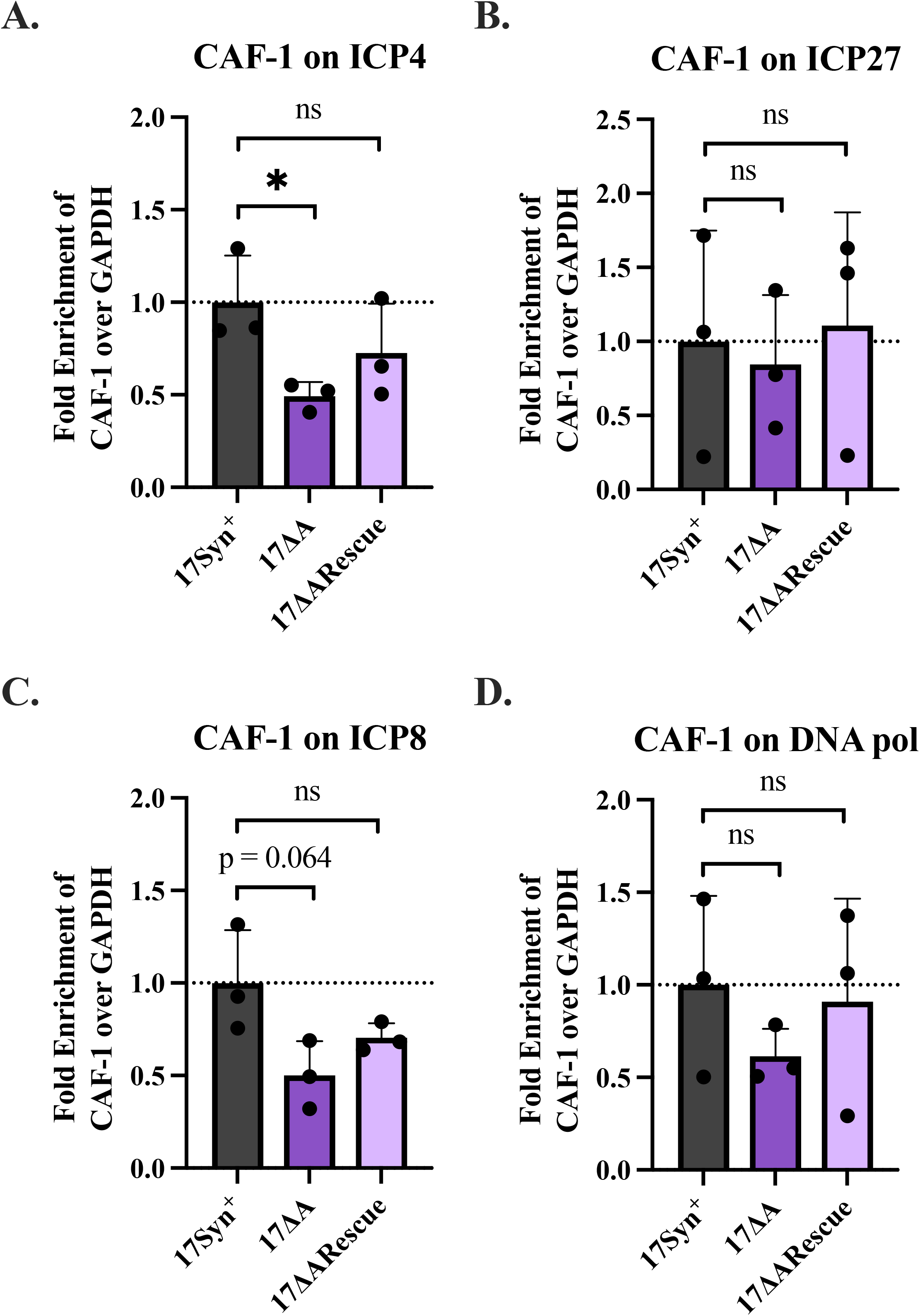
CAF-1 enrichment on viral lytic genes in 17syn+, 17ΔA, 17ΔARescue at 2 hpi. NHDF cells were infected with respective viruses at a MOI = 1, then ChIP-qPCR was performed to quantify CAF-1 enrichment on viral lytic genes and normalized to H3.1 enrichment on the cellular control gene, GAPDH. Fold enrichment is plotted against the 17syn+ sample (set to 1). Viral genes: **A)** ICP4, **B)** ICP27, **C)** ICP8, and **D)** HSV-1 DNA polymerase. N = 3. Unpaired two-tailed Student’s t-tests were performed to determine statistical significance between samples, means and standard deviations are plotted, * = p < 0.05, ns = not significant, p ≥ 0.05.

### CAF-1 depletion does not substantially alter H3.1 enrichment at viral lytic genes

The reduced CAF-1 occupancy observed following LTE deletion raised the possibility that CAF-1 contributed to LTE-dependent H3.1 enrichment. We therefore asked whether CAF-1 is required for H3.1 enrichment at viral lytic genes. To address this, we utilized siRNA to the CAF-1 subunit *CHAF1A*. As a control, total GAPDH RNA levels were measured for uninfected cells treated with the siScram and siCHAF1A **(Fig. S3A).** We then depleted the CHAF1A subunit of CAF-1 using siRNA and confirmed an approximately 80% reduction in CHAF1A protein levels compared with control siRNA-treated cells (**Fig. S3B**). Cells were then infected with WT HSV-1, and H3.1 enrichment at the viral loci examined above was quantified by ChIP-qPCR. Despite substantial depletion of CHAF1A, H3.1 enrichment at these viral loci was largely maintained and showed no significant reduction compared with control cells (**Fig. 7A-D; Fig. S4**). These findings indicate that H3.1 enrichment on the incoming HSV-1 genome was not solely dependent on CAF-1 under the conditions examined. Together with the reduced CAF-1 occupancy observed following LTE deletion, these results suggested that the LTE coordinated a locus-specific chromatin assembly process that is associated with both H3.1 enrichment and CAF-1 occupancy, while indicating that CAF-1 was not the sole determinant of H3.1 enrichment at these viral loci.

**Figure 7:**
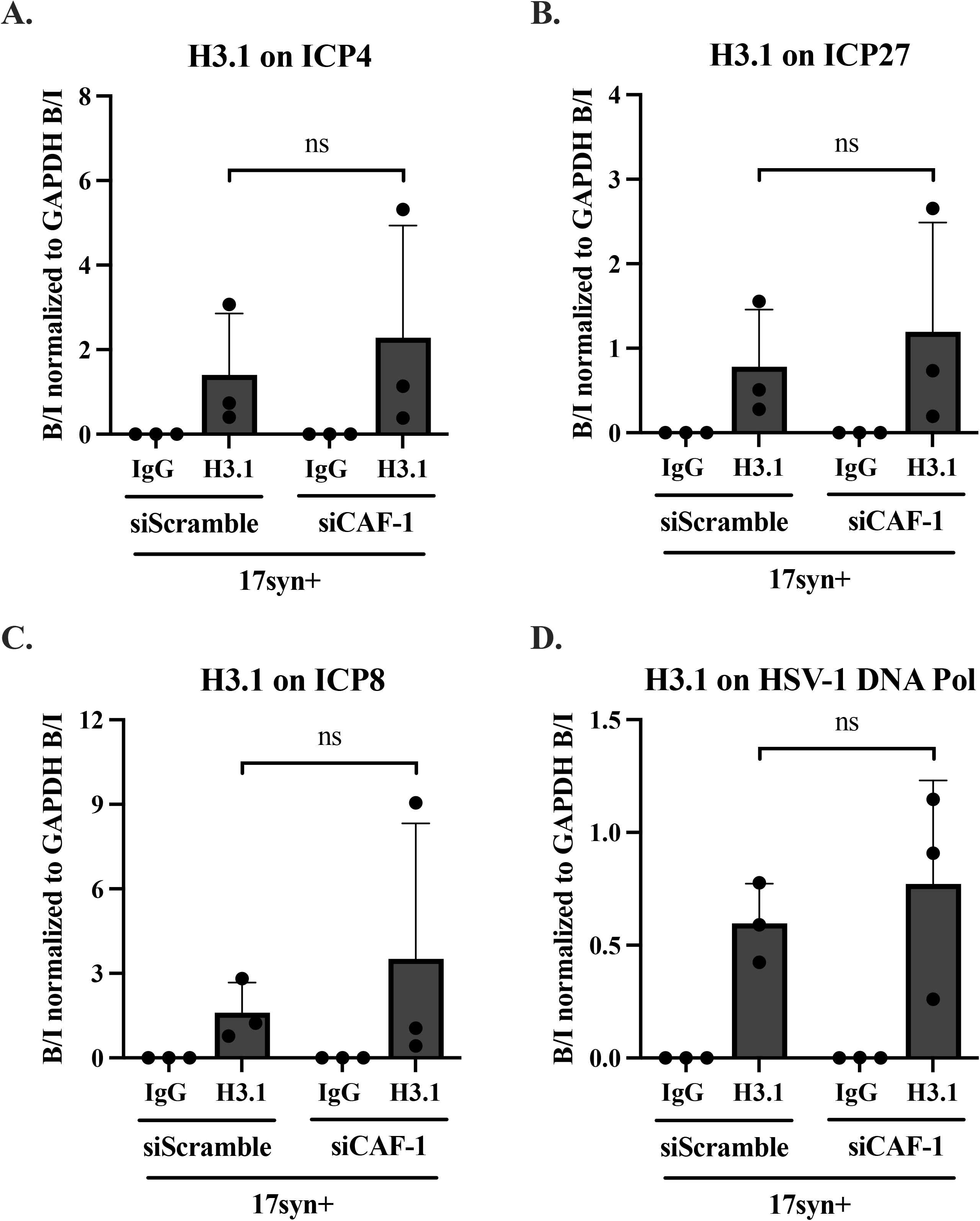
H3.1 enrichment on viral lytic genes is not altered by CAF-1 siRNA knockdown at 2 hpi in 17syn+. NHDF cells were transfected with either a non-targeting siRNA (siScramble) or siCAF-1 (against CHAF1A, the CAF-1 major subunit) for 48 hrs. Then, cells were infected with 17syn+ at a MOI =1. ChIP-qPCR was performed to quantify H3.1 enrichment on viral lytic genes and normalized to H3.1 enrichment on the cellular control gene, GAPDH. H3.1 enrichment is plotted against the non-specific binding control IgG (IgG values are the bound/input values of the respective viral gene). Viral genes: **A)** ICP4, **B)** ICP27, **C)** ICP8, and **D)** HSV-1 DNA polymerase. N = 3. Unpaired two-tailed Student’s t-tests were performed to determine statistical significance between samples, means and standard deviations are plotted, ns = not significant, p ≥ 0.05.

## Discussion

Gene regulation within mammalian cells is controlled by multiple factors, including dynamic changes in chromatin composition and organization. DNA viruses similarly exploit host transcriptional regulatory mechanisms to control viral gene expression. HSV-1 genomes rapidly associate with host histones following nuclear entry and changes in histone enrichment or modification contribute to the regulation of both lytic and latent infections of HSV-1. However, how specific viral regulatory elements influence the composition of chromatin assembled on the incoming viral genome remains poorly understood. Here, we identify a previously uncharacterized function of the HSV-1 LTE during lytic infection. We showed that a 310 bp region within the LTE contributed to locus-specific enrichment of the canonical H3.1 histone variant at selected viral lytic genes, including ICP4. This effect was distinct from the established enhancer activity of the LTE, was not accompanied by altered H3.3 enrichment at the viral loci examined and occurred independently of viral DNA replication. LTE deletion also reduced CAF-1 occupancy at loci with decreased H3.1 enrichment; however, depletion of CAF-1 did not substantially reduce H3.1 enrichment, indicating that CAF-1 was not the sole determinant of H3.1 enrichment on the incoming HSV-1 genome. Together, these findings suggest that the LTE coordinates a locus-specific chromatin environment during early lytic infection through mechanisms that influence H3.1 enrichment and CAF-1 occupancy.

Deletion of the 310-bp region in 17ΔA resulted in reduced H3.1 enrichment at ICP4 and ICP8, whereas enrichment at ICP27 and DNA polymerase remained unchanged. Thus, the effect of LTE deletion was neither global nor uniform across the viral genome but instead occurred at specific viral loci. Importantly, the phenotype was restored in the 17ΔA rescue virus, supporting a direct relationship between the deleted LTE sequence and H3.1 enrichment. Furthermore, deletion of the larger overlapping LTE region in 17Δ348, which also disrupted the established enhancer activity of the LTE, did not reproduce the H3.1 phenotype. These findings indicate that regulation of H3.1 enrichment is separable from the enhancer activity of the LTE and instead implicates sequence(s) within the 310-bp region deleted in 17ΔA.

The mechanisms by which this region of the LTE influenced H3.1 enrichment remain to be determined. The 17ΔA and 17Δ348 deletions overlap but differ in their non-overlapping sequences, providing a genetic framework for future identification of the sequence elements responsible for this activity. We predict that the deleted region contains binding sites for cellular or viral factors that influence chromatin assembly or histone chaperone occupancy on the viral genome. Determining which factors bind this region and whether their occupancy changes following LTE deletion so that we can define how a viral enhancer-associated sequence influences chromatin composition at distal viral loci are ongoing and will be reported in an additional manuscript. Further, the implications of this altered H3.1 occupancy and its impacts on efficient production of infectious progeny virions and how that altered occupancy affects the establishment of latency are also ongoing. It is completely possible that while we cannot detect differences in genome copies, the fitness of the progeny virions may be altered in the absence of the 310 bp LTE sequence.

The selective effect on H3.1 occupancy is particularly notable because H3.1 and H3.3 are incorporated into chromatin through distinct pathways and contribute to different chromatin environments. Although both variants are enriched on the HSV-1 genome during lytic infection, deletion of the LTE altered H3.1 but not H3.3 enrichment at the viral loci examined. These findings suggest that the LTE does not simply promote histone incorporation onto the viral genome but instead influences the composition of the histones that become associated with specific viral regions.

During lytic infection, HSV-1 executes a highly ordered temporal cascade of gene expression that is precipitated by the expression of five immediate early genes, including ICP4, a required gene for successful productive infection [35, 36]. ICP4 functions as a major transcriptional activator via binding to viral promoter regions and coordinating the formation of a pre-initiation complex to stimulate transcription [37, 38]. At the same time, ICP4 expression is subject to autoregulatory mechanisms that limit ICP4 expression and prevents excessive accumulation [39–41]. Controlling levels of ICP4 is particularly important for HSV-1, because elevated levels of ICP4 have been shown to be cytotoxic to the cell [42]. Thus, HSV-1 must express an appropriate level of ICP4 to initiate the viral temporal cascade, while at the same time preventing overaccumulation that can be detrimental to the host cell. Our findings raise the possibility that LTE-dependent H3.1 enrichment contributes to this regulation. The most pronounced chromatin phenotype was observed at ICP4, where deletion of the 310-bp LTE region reduced H3.1 enrichment without altering H3.3 enrichment. Consistent with a regulatory role for the LTE, 17ΔA also altered ICP4 expression, although the direction and magnitude of the transcriptional phenotype differed from those observed for other viral genes. At later times, ICP4 transcript abundance was increased in 17ΔA-infected cells, whereas ICP27, ICP8, and DNA polymerase transcripts remained reduced. The increased ICP4 expression in the context of reduced H3.1 enrichment is consistent with the possibility that LTE-dependent H3.1 enrichment contributes to a chromatin environment that constrains ICP4 transcription. However, the present experiments do not establish a direct causal relationship between H3.1 enrichment and ICP4 expression, and additional experiments are underway to determine whether altered H3.1 occupancy directly contributes to the transcriptional phenotype.

The observation that LTE deletion affects H3.1 enrichment by 2 hpi raises the question of how a replication-associated histone variant is incorporated onto the viral genome before substantial viral DNA replication. Consistent with previous studies, H3.1 is enriched on the HSV-1 genome during early lytic infection, while H3.3 enrichment is also detected during this period [7]. Our finding that inhibition of viral DNA replication did not eliminate the difference in H3.1 enrichment between WT and 17ΔA indicated that LTE-dependent H3.1 enrichment did not require viral genome replication. Instead, the LTE appears to influence chromatin assembly on the incoming viral genome before the onset of robust viral DNA synthesis.

We found that CAF-1 occupancy was reduced at ICP4 and ICP8 following deletion of the 310-bp LTE region, coincident with the reduction in H3.1 enrichment at these loci. This association suggests that the LTE influences the local chromatin assembly environment in a manner that affects both H3.1 and CAF-1 occupancy. However, CAF-1 depletion did not substantially reduce H3.1 enrichment at the viral loci examined, therefore the reduction in CAF-1 occupancy in the LTE deletion cannot be interpreted as a simple LTE–CAF-1–H3.1 pathway. Instead, our data indicate that CAF-1 may contribute to H3.1 enrichment while additional mechanisms can maintain H3.1 association with the viral genome when CAF-1 levels are reduced. This interpretation is consistent with previous studies demonstrating that HSV-1 can recruit multiple histone chaperones during initial genome chromatinization. For example, Cabral *et al.* reported that depletion of individual histone chaperones produced relatively modest effects on total H3 enrichment on the HSV-1 genome, whereas combined depletion of HIRA, Asf1, and ATRX produced a greater reduction in H3 enrichment [43, 44]. Their observation that CHAF1A depletion produced nonsignificant changes in H3 enrichment at ICP27 and ICP8 is consistent with our findings using H3.1-specific ChIP. Although their analysis examined total H3 rather than H3.1, H3.1 represents a substantial fraction of the canonical H3 pool in actively dividing cells [45]. Together, these studies support a model in which multiple histone chaperone pathways contribute to HSV-1 chromatin assembly and may provide partially redundant mechanisms for maintaining histone enrichment on the incoming genome.

The findings presented here expand the functional repertoire of the HSV-1 LTE beyond its established roles in latency and reactivation. Rather than functioning solely as a transcriptional enhancer, the LTE contains sequence information that influences the chromatin environment established on the viral genome during early lytic infection. Our findings also raise the possibility that individual viral regulatory elements can shape the chromatin landscape of the HSV-1 genome in a locus-and histone-variant-specific manner. Such mechanisms could allow HSV-1 to establish distinct chromatin environments across different regions of its genome during the earliest stages of infection, thereby contributing to the ordered expression of viral genes. Whether the LTE influences chromatin directly through binding of specific cellular factors or indirectly by altering higher-order viral genome organization remains to be determined.

## Methods and Materials

### Cells and Viruses

Normal Human Dermal Fibroblast (NHDF) (ATCC PCS-201-012) cells were cultured in Dulbecco’s Modified Eagle’s Medium (DMEM; Corning 15013) supplemented with 15% fetal bovine serum (FBS; Corning 35011CV) and 1% antibiotic-antimycotic solution (Corning 30-004-CI). The cells were maintained in a humidified incubator at 37°C and 5% CO_2_. Cells were regularly passaged to keep confluency below 80%. All experiments were conducted with cells from below passage 20.

The wild type HSV-1 strain, 17*Syn*+ (GenBank NC_001806) was gifted from J. Stevens. The recombinant LTE deletion viruses: 17ΔA (deletion at n.t.119,197-119,507), 17ΔARescue, 17Δ348 (deletion at n.t.119,007-119,355), 17Δ348Rescue strains were gifted from D. Bloom (University of Florida). All recombinants were derived from the parental 17*Syn*+ strain. These viruses were propagated in Vero cells (ATCC CCL-81) and titered in the Neumann lab.

### *In vitro* infections

NHDF cells were seeded in 15% DMEM into 6-well plates or T-25 flasks and grown until 80-90% confluency. Cells were infected with WT or a recombinant virus at MOI of 1 in 1% DMEM. Infected cells were rocked at 4°C for 1 hr then incubated at 37°C for 1 hr, subsequently the media was exchanged for virus-free 1% DMEM. For experiments assessing protein enrichment in the absence of viral replication, an HSV-1 replication inhibitor was used. Infection was performed as described, except all media was supplemented with 200 μg/mL Phosphonoacetic acid (PAA; ThermoScientific A12117.09) beginning at time of infection until sample collection was performed.

### Viral genome copies and plaque assay

NHDF cells were infected with WT or 17ΔA at MOI of 0.1. Infected cells were collected at 8, 24, 48, and 72 hpi in SDS-lysis buffer and incubated at 65°C overnight. Total DNA was isolated by phenol-chloroform extraction. Viral genome copy numbers were determined using qPCR with primers and probes specific for the HSV-1 DNA polymerase and normalized to the host control gene GAPDH. For plaque assays, supernatant was collected from infected cells at 8, 24, 48, and 72 hpi. The samples were serial diluted and plated onto 100% confluent Vero cells in 24-well plates. The plates were incubated at 37°C and 5% CO_2_ for 48 hrs. Media was removed and cells were fixed and stained by adding 500 µL of BBL Gram Crystal Violet (BD B12525) and incubated at 25°C overnight. Plates were rinsed with water and plaque forming units were quantitated.

### Chromatin Immunoprecipitation (ChIP) and quantitative PCR (qPCR)

ChIP samples were collected and washed with ice-cold 1X Phosphate-Buffered Saline (PBS) with protease inhibitors (2mg/mL Aprotinin, 5mg/mL Leupeptin, 10mM PMSF). At room temp, cells were crosslinked using a final concentration of 1% formaldehyde (RICCA RSF0010-500A) and subsequently the chemical reaction was quenched using a final concentration of 125mM glycine (Fisher chemical G46-1). Then, samples were washed with PBS containing protease inhibitors three times and resuspended in ChIP SDS-lysis buffer containing protease inhibitors and incubated for 30 mins on ice. Samples were sonicated to shear the chromatin into 300-800 bp fragments (QSonica Q800R3 Sonicator) as follows: 50% amplitude, 30s on, 50s off for 24 cycles while samples are kept under 4°C during sonication. Sheared samples were centrifuged at 4°C for 10 mins at 13,500 rpm and the supernatant transferred to a new tube and the pellet discarded. The supernatant was diluted 10:1 using the ChIP Dilution Buffer and separated into fractions for IgG, Antibody bound, and Input samples. Each fraction was incubated with its respective antibody overnight at 4°C while rocking. Antibodies used include: IgG (EMD Millipore CS200581), H3.1/H3.1 (EMD Millipore ABE154), H3.3 (Abcam AB176840) and CAF-1 p150 (Active Motif 39997). Chromatin bound to antibodies were captured via a magnetic bead pulldown. Protein G magnetic beads (Cell Signaling 9006S) were added to each fraction (30 μL per 1 mL sample) and rocked at 4°C for 2 hrs. ChIP complexes were washed using sequential rinses of 150 mM Low Salt Immune Complex Wash, 500 mM High Salt Immune Complex Wash, 0.25 LiCl Immune Complex Wash, TE Buffer, then eluted in 950 μL Elution Buffer. Crosslinking was reversed by adding 5 M NaCl (final concentration 200 mM) to eluted and input samples while incubating at 65°C for 4 hrs. Then all RNA and protein are degraded by incubating samples with 2 μL/mL RNAse A (ThermoScientific EN0531) at 37°C for 30 mins, then with 2 μL/mL proteinase K (ThermoScientific EO0491) at 65°C for 2 hrs. DNA was purified by column purification using the QIAquick Nucleotide Removal kit (Qiagen 28306) following the manufacturer’s protocol and eluted into 80 μL elution buffer. qPCRs were performed using TaqMan™ Fast Universal PCR Master Mix (2×), no AmpErase™ UNG (Applied biosystems 4352042) on an Agilent AriaMx Real-Time PCR machine. The following PCR cycling conditions were used: 95 °C for 10 min and then 95 °C for 15 sec, followed by 60°C for 1 min (x40 cycles).

### Small interfering RNA (siRNA) transfections

NHDF cells were plated in 6-well plates at 2.5 x 10^5^ cells per well in 15% DMEM and grown for 48 hrs at 37°C and 5% CO2. siRNAs were transfected in Opti-MEM I (Gibco 31985-070) and 7.5 μL Lipofectamine RNAiMAX Reagent (Invitrogen 13778-150) per well, following the manufacturer’s protocol. A non-targeting siRNA (siScramble; Dharmacon D-001810-01-05) and siRNA against a subunit of CAF-1 (siCHAF1A; Dharmacon J-019938-09-0002) were transfected at 40 pmol per well for 48 hrs. Knockdown of CHAF1A was verified by measuring CHAF1A transcript levels and >80% knockdown was achieved (Figure S3). Then, transfected cells were infected with 17syn+ at an MOI of 1, as described above.

### Quantitative reverse transcriptase PCR (qRT-PCR)

qRT-PCR was performed to compare gene expression between WT and LTE deletion viruses. NHDF cells were infected with WT or a recombinant virus at an MOI of 1 in 6-well plates. Cells were harvested in 500 µL of TRIzol reagent (Ambion 15596026) per well at 2, 6, and 24 hpi. Total RNA was extracted according to the manufacturer’s protocol. qRT-PCR reactions were set up according to the manufacturer’s protocol using the SuperScript III Platinum One-Step qRT-PCR System (Invitrogen 11732088) and Taqman primer probe mixtures (Table 1). The following PCR cycling conditions were used: 50°C for 15 min, 95°C for 5 min, and then 95°C for 15 s followed by 60°C for 30 sec (x40 cycles). Purified NHDF or 17syn+ RNA was used to generate a standard curve for each qRT-PCR plate and sample copy numbers were determined based on this linear range.

### Figure Generation and Statistical analyses

BioRender and PowerPoint were used to generate Figure 1. GraphPad Prism, version 11.1.0 for Mac, was used to generate all bar graphs and perform statistical analyses using Student T-Tests.

## Supporting information

combined supplemental figures

## Acknowledgements

This work was supported by the grants NIH/NIAID R01AI19006 (DMN), NIH/NEI 2T32EY027721 (MAS), the Core Grant for Vision Research from the NIH to the University of Wisconsin-Madison (P30 EY016665), the McPherson Eye Research Institute Grant Summit Program and an unrestricted grant from Research to Prevent Blindness (Department of Ophthalmology, University of Wisconsin). The funders had no role in study design, data collection and analysis, decision to publish, or preparation of the manuscript.

## References

1. Austin A, Lietman T, Rose-Nussbaumer J. Update on the Management of Infectious Keratitis. Ophthalmology. 2017;124(11):1678–89. Epub 2017/09/25. doi: 10.1016/j.ophtha.2017.05.012. PubMed PMID: 28942073; PubMed Central PMCID: PMCPMC5710829.

2. Kolb AW, Ferguson SA, Larsen IV, Brandt CR. Disease parameters following ocular herpes simplex virus type 1 infection are similar in male and female BALB/C mice. PLoS One. 2023;18(6):e0287194. Epub 20230615. doi: 10.1371/journal.pone.0287194. PubMed PMID: 37319284; PubMed Central PMCID: PMCPMC10270577.

3. Zhu S, Viejo-Borbolla A. Pathogenesis and virulence of herpes simplex virus. Virulence. 2021;12(1):2670–702. doi: 10.1080/21505594.2021.1982373. PubMed PMID: 34676800; PubMed Central PMCID: PMCPMC8923070.

4. Kaufman HE. Does everyone have herpes? Cornea. 2004;23(2):111–2. doi: 10.1097/00003226-200403000-00001. PubMed PMID: 15075877.

5. Lacasse JJ, Schang LM. Herpes simplex virus 1 DNA is in unstable nucleosomes throughout the lytic infection cycle, and the instability of the nucleosomes is independent of DNA replication. J Virol. 2012;86(20):11287–300. Epub 20120808. doi: 10.1128/JVI.01468-12. PubMed PMID: 22875975; PubMed Central PMCID: PMCPMC3457126.

6. Kent JR, Zeng PY, Atanasiu D, Gardner J, Fraser NW, Berger SL. During lytic infection herpes simplex virus type 1 is associated with histones bearing modifications that correlate with active transcription. J Virol. 2004;78(18):10178–86. doi: 10.1128/JVI.78.18.10178-10186.2004. PubMed PMID: 15331750; PubMed Central PMCID: PMCPMC514973.

7. Placek BJ, Huang J, Kent JR, Dorsey J, Rice L, Fraser NW, et al. The histone variant H3.3 regulates gene expression during lytic infection with herpes simplex virus type 1. J Virol. 2009;83(3):1416–21. Epub 20081112. doi: 10.1128/JVI.01276-08. PubMed PMID: 19004946; PubMed Central PMCID: PMCPMC2620911.

8. Conn KL, Schang LM. Chromatin dynamics during lytic infection with herpes simplex virus 1. Viruses. 2013;5(7):1758–86. Epub 20130716. doi: 10.3390/v5071758. PubMed PMID: 23863878; PubMed Central PMCID: PMCPMC3738960.

9. Conn KL, Hendzel MJ, Schang LM. The differential mobilization of histones H3.1 and H3.3 by herpes simplex virus 1 relates histone dynamics to the assembly of viral chromatin. PLoS Pathog. 2013;9(10):e1003695. Epub 20131010. doi: 10.1371/journal.ppat.1003695. PubMed PMID: 24130491; PubMed Central PMCID: PMCPMC3795045.

10. Conn KL, Hendzel MJ, Schang LM. Core histones H2B and H4 are mobilized during infection with herpes simplex virus 1. J Virol. 2011;85(24):13234–52. Epub 20111012. doi: 10.1128/JVI.06038-11. PubMed PMID: 21994445; PubMed Central PMCID: PMCPMC3233158.

11. Hegazy YA, Dhahri H, El Osmani N, George S, Chandler DP, Fondufe-Mittendorf YN. Histone variants: The bricks that fit differently. J Biol Chem. 2025;301(1):108048. Epub 20241204. doi: 10.1016/j.jbc.2024.108048. PubMed PMID: 39638247; PubMed Central PMCID: PMCPMC11742582.

12. Talbert PB, Henikoff S. Histone variants at a glance. J Cell Sci. 2021;134(6). Epub 20210326. doi: 10.1242/jcs.244749. PubMed PMID: 33771851; PubMed Central PMCID: PMCPMC8015243.

13. Tagami H, Ray-Gallet D, Almouzni G, Nakatani Y. Histone H3.1 and H3.3 complexes mediate nucleosome assembly pathways dependent or independent of DNA synthesis. Cell. 2004;116(1):51–61. doi: 10.1016/s0092-8674(03)01064-x. PubMed PMID: 14718166.

14. Chen P, Zhao J, Wang Y, Wang M, Long H, Liang D, et al. H3.3 actively marks enhancers and primes gene transcription via opening higher-ordered chromatin. Genes Dev. 2013;27(19):2109–24. Epub 20130924. doi: 10.1101/gad.222174.113. PubMed PMID: 24065740; PubMed Central PMCID: PMCPMC3850095.

15. Jin C, Felsenfeld G. Nucleosome stability mediated by histone variants H3.3 and H2A.Z. Genes Dev. 2007;21(12):1519–29. doi: 10.1101/gad.1547707. PubMed PMID: 17575053; PubMed Central PMCID: PMCPMC1891429.

16. Goldberg AD, Banaszynski LA, Noh KM, Lewis PW, Elsaesser SJ, Stadler S, et al. Distinct factors control histone variant H3.3 localization at specific genomic regions. Cell. 2010;140(5):678–91. doi: 10.1016/j.cell.2010.01.003. PubMed PMID: 20211137; PubMed Central PMCID: PMCPMC2885838.

17. García-González E, Escamilla-Del-Arenal M, Arzate-Mejía R, Recillas-Targa F. Chromatin remodeling effects on enhancer activity. Cell Mol Life Sci. 2016;73(15):2897–910. Epub 20160330. doi: 10.1007/s00018-016-2184-3. PubMed PMID: 27026300; PubMed Central PMCID: PMCPMC11108574.

18. Rao SS, Huntley MH, Durand NC, Stamenova EK, Bochkov ID, Robinson JT, et al. A 3D map of the human genome at kilobase resolution reveals principles of chromatin looping. Cell. 2014;159(7):1665–80. Epub 20141211. doi: 10.1016/j.cell.2014.11.021. PubMed PMID: 25497547; PubMed Central PMCID: PMCPMC5635824.

19. Tang F, Yang Z, Tan Y, Li Y. Super-enhancer function and its application in cancer targeted therapy. NPJ Precis Oncol. 2020;4:2. Epub 20200212. doi: 10.1038/s41698-020-0108-z. PubMed PMID: 32128448; PubMed Central PMCID: PMCPMC7016125.

20. Drolet BS, Perng GC, Villosis RJ, Slanina SM, Nesburn AB, Wechsler SL. Expression of the first 811 nucleotides of the herpes simplex virus type 1 latency-associated transcript (LAT) partially restores wild-type spontaneous reactivation to a LAT-null mutant. Virology. 1999;253(1):96–106. doi: 10.1006/viro.1998.9492. PubMed PMID: 9887322.

21. Perng GC, Slanina SM, Yukht A, Drolet BS, Keleher W, Ghiasi H, et al. A herpes simplex virus type 1 latency-associated transcript mutant with increased virulence and reduced spontaneous reactivation. J Virol. 1999;73(2):920–9. doi: 10.1128/JVI.73.2.920-929.1999. PubMed PMID: 9882292; PubMed Central PMCID: PMCPMC103911.

22. Perng GC, Dunkel EC, Geary PA, Slanina SM, Ghiasi H, Kaiwar R, et al. The latency-associated transcript gene of herpes simplex virus type 1 (HSV-1) is required for efficient in vivo spontaneous reactivation of HSV-1 from latency. J Virol. 1994;68(12):8045–55. doi: 10.1128/JVI.68.12.8045-8055.1994. PubMed PMID: 7966594; PubMed Central PMCID: PMCPMC237268.

23. Ahmed M, Lock M, Miller CG, Fraser NW. Regions of the herpes simplex virus type 1 latency-associated transcript that protect cells from apoptosis in vitro and protect neuronal cells in vivo. J Virol. 2002;76(2):717–29. doi: 10.1128/jvi.76.2.717-729.2002. PubMed PMID: 11752162; PubMed Central PMCID: PMCPMC136840.

24. Perng GC, Jones C, Ciacci-Zanella J, Stone M, Henderson G, Yukht A, et al. Virus-induced neuronal apoptosis blocked by the herpes simplex virus latency-associated transcript. Science. 2000;287(5457):1500–3. doi: 10.1126/science.287.5457.1500. PubMed PMID: 10688801.

25. Wang QY, Zhou C, Johnson KE, Colgrove RC, Coen DM, Knipe DM. Herpesviral latency-associated transcript gene promotes assembly of heterochromatin on viral lytic-gene promoters in latent infection. Proc Natl Acad Sci U S A. 2005;102(44):16055–9. Epub 20051024. doi: 10.1073/pnas.0505850102. PubMed PMID: 16247011; PubMed Central PMCID: PMCPMC1266038.

26. Cliffe AR, Garber DA, Knipe DM. Transcription of the herpes simplex virus latency-associated transcript promotes the formation of facultative heterochromatin on lytic promoters. J Virol. 2009;83(16):8182–90. Epub 2009/06/12. doi: 10.1128/JVI.00712-09. PubMed PMID: 19515781; PubMed Central PMCID: PMCPMC2715743.

27. Bloom DC, Hill JM, Devi-Rao G, Wagner EK, Feldman LT, Stevens JG. A 348-base-pair region in the latency-associated transcript facilitates herpes simplex virus type 1 reactivation. J Virol. 1996;70(4):2449–59. doi: 10.1128/JVI.70.4.2449-2459.1996. PubMed PMID: 8642650; PubMed Central PMCID: PMCPMC190088.

28. Berthomme H, Thomas J, Texier P, Epstein A, Feldman LT. Enhancer and long-term expression functions of herpes simplex virus type 1 latency-associated promoter are both located in the same region. J Virol. 2001;75(9):4386–93. doi: 10.1128/JVI.75.9.4386-4393.2001. PubMed PMID: 11287587; PubMed Central PMCID: PMCPMC114183.

29. Berthomme H, Lokensgard J, Yang L, Margolis T, Feldman LT. Evidence for a bidirectional element located downstream from the herpes simplex virus type 1 latency-associated promoter that increases its activity during latency. J Virol. 2000;74(8):3613–22. doi: 10.1128/jvi.74.8.3613-3622.2000. PubMed PMID: 10729137; PubMed Central PMCID: PMCPMC111871.

30. Alyssa Richman SK, Terri Edwards, Joseph Boyd, Princess Rodriguez, Michael Mariani, Mason A. Shipley https://orcid.org/0000-0001-8317-3166, Kayley A. Manuel https://orcid.org/0009-0005-5145-4015, Ziyun A. Ye https://orcid.org/0009-0007-5819-6957, David C. Bloom, Seth Frietze https://orcid.org/0000-0003-4058-3661, Donna M. Neumann https://orcid.org/0000-0002-8115-2254. Multiple long-range cis interactions generate CTCF insulator-dependent viral chromatin domains in quiescent HSV-1 genomes.

31. Ye ZA, Shipley MA, Singh P, Neumann DM. In HSV-1, the LAT Enhancer Drives Pre-IE VP16 Transcription to Initiate Reactivation. bioRxiv. 2026. Epub 20260106. doi: 10.64898/2026.01.06.697999. PubMed PMID: 41542627; PubMed Central PMCID: PMCPMC12803242.

32. Overby LR, Robishaw EE, Schleicher JB, Rueter A, Shipkowitz NL, Mao JC. Inhibition of herpes simplex virus replication by phosphonoacetic acid. Antimicrob Agents Chemother. 1974;6(3):360–5. doi: 10.1128/AAC.6.3.360. PubMed PMID: 15830487; PubMed Central PMCID: PMCPMC444651.

33. Crumpacker CS. Mechanism of action of foscarnet against viral polymerases. Am J Med. 1992;92(2A):3S–7S. doi: 10.1016/0002-9343(92)90329-a. PubMed PMID: 1371038.

34. Smith S, Stillman B. Purification and characterization of CAF-I, a human cell factor required for chromatin assembly during DNA replication in vitro. Cell. 1989;58(1):15–25. doi: 10.1016/0092-8674(89)90398-x. PubMed PMID: 2546672.

35. DeLuca NA, Schaffer PA. Activation of immediate-early, early, and late promoters by temperature-sensitive and wild-type forms of herpes simplex virus type 1 protein ICP4. Mol Cell Biol. 1985;5(8):1997–2008. doi: 10.1128/mcb.5.8.1997-2008.1985. PubMed PMID: 3018543; PubMed Central PMCID: PMCPMC366918.

36. DeLuca NA, McCarthy AM, Schaffer PA. Isolation and characterization of deletion mutants of herpes simplex virus type 1 in the gene encoding immediate-early regulatory protein ICP4. J Virol. 1985;56(2):558–70. doi: 10.1128/JVI.56.2.558-570.1985. PubMed PMID: 2997476; PubMed Central PMCID: PMCPMC252613.

37. Grondin B, DeLuca N. Herpes simplex virus type 1 ICP4 promotes transcription preinitiation complex formation by enhancing the binding of TFIID to DNA. J Virol. 2000;74(24):11504–10. doi: 10.1128/jvi.74.24.11504-11510.2000. PubMed PMID: 11090147; PubMed Central PMCID: PMCPMC112430.

38. Smith CA, Bates P, Rivera-Gonzalez R, Gu B, DeLuca NA. ICP4, the major transcriptional regulatory protein of herpes simplex virus type 1, forms a tripartite complex with TATA-binding protein and TFIIB. J Virol. 1993;67(8):4676–87. doi: 10.1128/JVI.67.8.4676-4687.1993. PubMed PMID: 8392607; PubMed Central PMCID: PMCPMC237853.

39. Godowski PJ, Knipe DM. Transcriptional control of herpesvirus gene expression: gene functions required for positive and negative regulation. Proc Natl Acad Sci U S A. 1986;83(2):256–60. doi: 10.1073/pnas.83.2.256. PubMed PMID: 3001729; PubMed Central PMCID: PMCPMC322836.

40. Gu B, Rivera-Gonzalez R, Smith CA, DeLuca NA. Herpes simplex virus infected cell polypeptide 4 preferentially represses Sp1-activated over basal transcription from its own promoter. Proc Natl Acad Sci U S A. 1993;90(20):9528–32. doi: 10.1073/pnas.90.20.9528. PubMed PMID: 8415735; PubMed Central PMCID: PMCPMC47602.

41. Chaturvedi S, Engel R, Weinberger L. The HSV-1 ICP4 Transcriptional Auto-Repression Circuit Functions as a Transcriptional “Accelerator” Circuit. Front Cell Infect Microbiol. 2020;10:265. Epub 20200624. doi: 10.3389/fcimb.2020.00265. PubMed PMID: 32670890; PubMed Central PMCID: PMCPMC7326776.

42. Krisky DM, Wolfe D, Goins WF, Marconi PC, Ramakrishnan R, Mata M, et al. Deletion of multiple immediate-early genes from herpes simplex virus reduces cytotoxicity and permits long-term gene expression in neurons. Gene Ther. 1998;5(12):1593–603. doi: 10.1038/sj.gt.3300766. PubMed PMID: 10023438.

43. Cabral JM, Cushman CH, Sodroski CN, Knipe DM. ATRX limits the accessibility of histone H3-occupied HSV genomes during lytic infection. PLoS Pathog. 2021;17(4):e1009567. Epub 20210428. doi: 10.1371/journal.ppat.1009567. PubMed PMID: 33909709; PubMed Central PMCID: PMCPMC8109836.

44. Latreille D, Bluy L, Benkirane M, Kiernan RE. Identification of histone 3 variant 2 interacting factors. Nucleic Acids Res. 2014;42(6):3542–50. Epub 20140106. doi: 10.1093/nar/gkt1355. PubMed PMID: 24393775; PubMed Central PMCID: PMCPMC3973350.

45. Scott WA, Campos EI. Interactions With Histone H3 & Tools to Study Them. Front Cell Dev Biol. 2020;8:701. Epub 20200731. doi: 10.3389/fcell.2020.00701. PubMed PMID: 32850821; PubMed Central PMCID: PMCPMC7411163.

