## Supplementary material for "The HSV-1 LAT enhancer regulates locus-specific H3.1 enrichment during early lytic infection": combined supplemental figures

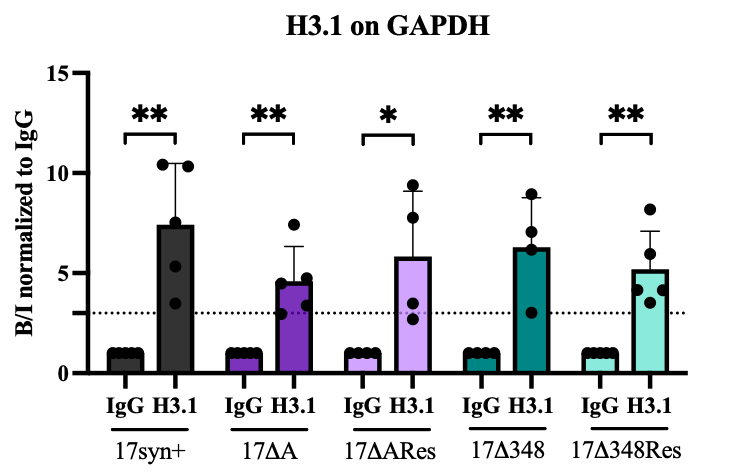

**Figure S1: H3.1 is significantly enriched on the cellular control gene GAPDH in 17syn+, 17ΔA, 17ΔARescue, 17Δ348, and 17Δ348Rescue at 2 hpi.**

NHDF cells were infected with respective viruses at a MOI = 1, then ChIP-qPCR was performed to quantify H3.1 enrichment compared to the non-specific binding control IgG. All H3.1 sample values were normalized to IgG (set to 1) to show the fold enrichment. The dotted line denotes a 3-fold enrichment of H3.1 compared to IgG. N = 4-5. Unpaired two-tailed Student’s t-tests were performed to determine statistical significance between samples, means and standard deviations are plotted, * = p < 0.05, ** = p < 0.01.

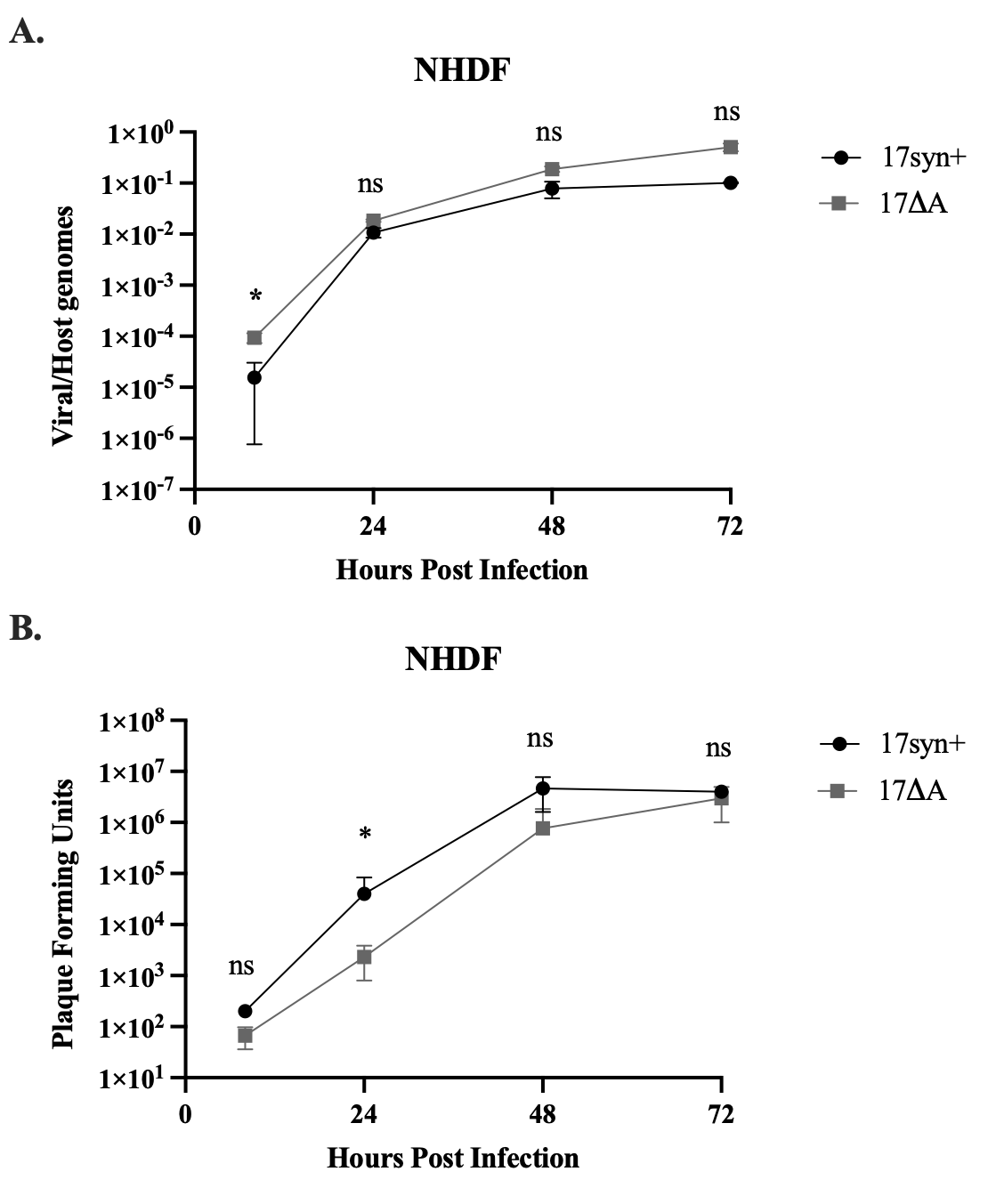

**Figure S2: 17ΔA exhibits no significant defect in viral replication or formation of infectious particles compared to 17syn+.**

NHDF cells were infected with a MOI = 0.1 and samples were collected at 8, 24, 48, and 72 hpi. **A)** DNA extraction was performed and qPCR was performed to quantitate human GAPDH and HSV-1 DNA polymerase, data plotted represent the fold change of DNA polymerase/GAPDH. **B)** Supernatant were collected to perform plaque assays to quantitate the amount of PFUs. N = 3. Unpaired two-tailed Student’s t-tests were performed to determine statistical significance between samples, means and standard deviations are plotted, * = p < 0.05, ns = not significant, p ≥ 0.05.

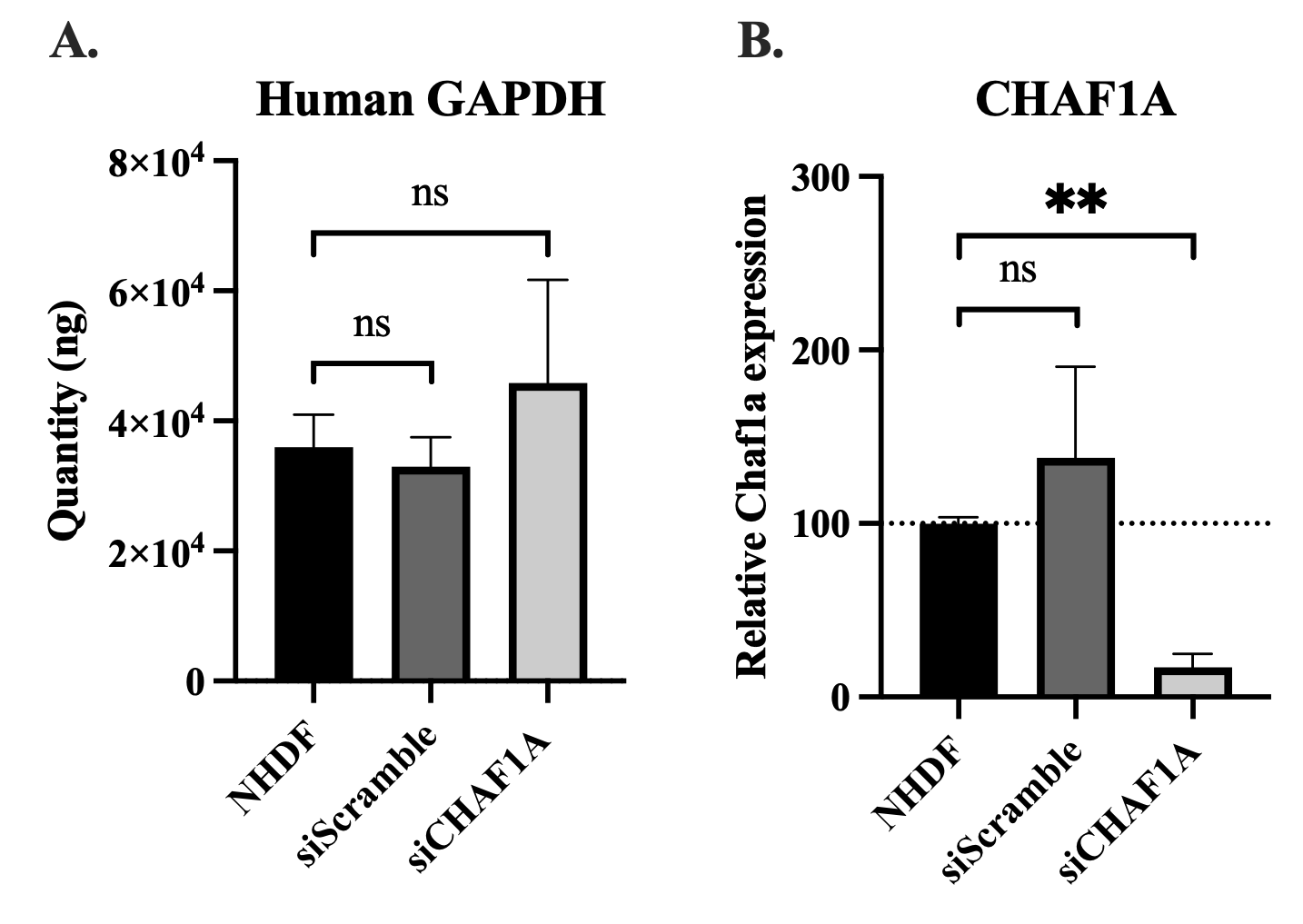

**Figure S3: siRNA knockdown of CAF-1 does not alter global cellular gene expression in uninfected NHDF cells.**

Uninfected NHDF cells underwent a 48-hr transfection with RNAi transfection reagents and respective siRNAs. Samples were collected and RNA was isolated and purified for use in qRT-PCR. **A)** Human GAPDH total RNA expression. **B)** Chaf1a (CAF-1 major subunit) relative expression normalized to human GAPDH expression. NHDF = Cells + RNAi transfection reagents, siScramble = non-targeting siRNA (negative control), siChaf1a = siRNA against the CAF-1 p150 subunit. N = 3. Unpaired two-tailed Student’s t-tests were performed to determine statistical significance between samples, means and standard deviations are plotted, ** = p < 0.01, ns = not significant, p ≥ 0.05.

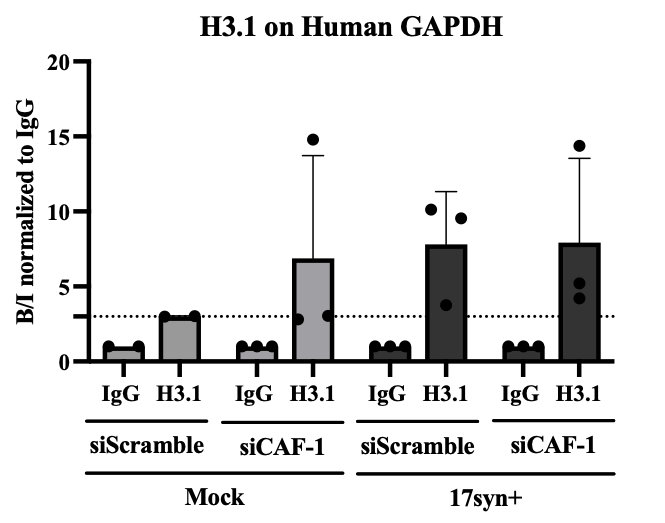

**Figure S4: H3.1 enrichment is not altered by siRNA knockdown of CAF-1 on the cellular host gene, GAPDH.**

Uninfected NHDF cells were transfected with respective siRNAS for 48 hrs (siScramble is the non-targeting siRNA and siCAF-1 targets Chaf1a), then samples were either collected (Mock) or cells were infected with 17syn+ at a MOI = 1. ChIP-qPCR was performed to quantify H3.1 enrichment compared to the non-specific binding control IgG. All H3.1 sample values were normalized to IgG (set to 1) to show the fold enrichment. The dotted line denotes a 3-fold enrichment of H3.1 compared to IgG. N = 3. Means and standard deviations are plotted.

| **Primer** | **Company** | **Catalog Number** |
| --- | --- | --- |
| Human GAPDH | TaqMan | Hs03929097_gl |
| ICP4 | TaqMan | APPRY2F |
| ICP8 | TaqMan | APKCHCN |
| ICP27 | TaqMan | APH6NRR |
| HSV-1 DNA Polymerase | TaqMan | AP7D4MW |
| VP16 | TaqMan | AIQI074 |

**Table S1: Primers and probes used in qPCR and RT-qPCR analyses.**
